# Hv1 channel supports insulin secretion in pancreatic *β* cells through calcium entry, depolarization and intracellular pH regulation

**DOI:** 10.1101/097816

**Authors:** Huimin Pang, Xudong Wang, Wang Xi, Qing Zhao, Shangrong Zhang, Jiwei Qin, Jili Lv, Yongzhe Che, Weiyan Zuo, Shu Jie Li

## Abstract

Here, we demonstrate that the voltage-gated proton channel Hv1 represents a regulatory mechanism for insulin secretion of pancreatic islet *β* cell. *In vivo*, Hv1-deficient mice display hyperglycemia and glucose intolerance due to reduced insulin secretion, but normal peripheral insulin sensitivity. *In vitro*, islets of Hv1-deficient and heterozygous mice, INS-1 (832/13) cells with siRNA-mediated knockdown of Hv1 exhibit a marked defect in glucose- and K^+^-induced insulin secretion. Hv1 deficiency decreases both insulin and proinsulin contents, and limits glucose-induced Ca^2+^ entry and membrane depolarization. Furthermore, loss of Hv1 increases insulin-containing granular pH and decreases cytosolic pH. In addition, histologic studies show a decrease in *β* cell mass in islets of Hv1-deficient mice. Collectively, our results indicate that Hv1 supports insulin secretion in the *β* cell by calcium entry, membrane depolarization and intracellular pH regulation.

**SIGNIFICANCE STATEMENT:** The voltage-gated proton channel Hv1 is highly expressed in insulin-containing granules in pancreatic β cells. Hv1 supports insulin secretion in the *β* cell by calcium entry, membrane depolarization and regulation of intragranular and cytosolic pH, which represents a regulatory mechanism for insulin secretion of pancreatic islet *β* cell. Our research demonstrates that Hv1 expressed in *β* cell is required for insulin secretion and maintains glucose homeostasis, and reveals a significant role for the proton channel in the modulation of pancreatic *β* cell function.

## INTRODUCTION

Insulin secretion by pancreatic *β* cells is precisely regulated by glucose homeostasis. The glucose metabolism by the pancreatic *β* cells accompanies proton generation, which proposes a mechanism of intracellular pH-regulation behind insulin release stimulated by the sugar (1). Manipulating intracellular as well as extracellular pH could affect the insulin secretory process, which is associated with changes in membrane potential, ionic fluxes, and insulin release (1-3). Even though some studies showed that glucose had no effect on intracellular pH (4,5), some reports showed that glucose induced cytosolic pH increase (6-8). Meanwhile, it is reported that intracellular alkalization inhibits insulin secretion from cells (3,9), acidification stimulates insulin release (10). Some studies proposed that Na^+^/H^+^ and Cl^-^/HCO_3_^-^ exchangers enable *β* cells to effectively buffer the acid load generated by glucose metabolism (6). However, either a decrease or an increase in pH_c_ is important for glucose-induced insulin secretion via the triggering or the amplifying pathways.

The pH of insulin-containing granules is between 5 and 6, which is thought to permit sequential action of pH-dependent prohormone convertases in the proteolytic processing of proinsulin and favor storage of insoluble Zn^2+^-insulin hexamers (11,12). The results that glucose induces pH changes in insulin-containing granules are contradictory. Glucose slightly increased pH of secretary vesicles in normal mouse islets (13) but acidified insulin granules in RIN insulinoma cells (14). Simultaneously, other study showed that cytosolic ATP promoted granule acidification (15).

The voltage-gated proton channel Hv1 is extremely selective for protons and has no detectable permeability to other cations (16,17). Hv1 channel is activated at depolarizing voltages, sensitive to the membrane pH gradient, H^+^-selective, Zn^2+^- and temperature-sensitive (16,17). In our previous study, we have identified that Hv1 is present in human and rodent pancreatic islet *β* cells, as well as *β* cell lines (18). However, the regulatory mechanism of Hv1 for insulin secretion of pancreatic islet *β* cell is not known. In present study, we have discovered a regulatory mechanism for the proton channel Hv1 in the modulation of *β* cell insulin secretory function. Our *in vitro* and *in vivo* studies demonstrate that Hv1 is required for insulin secretion in the *β* cell.

## RESULTS

### Hv1-deficient mice exhibit hyperglycaemia and impaired glucose tolerance due to reduced insulin secretion

To assess the effect of Hv1 knockout on glucose homeostasis, glucose levels were measured in 4 month-old mice in fasted state. The body weight curves of the control (WT, Hv1^+/+^), heterozygous (Hv1^+/−^) and homozygous (KO, Hv1^−/−^) littermates were almost similar (Fig. 1A), but the blood glucose levels in fasted state were markedly higher in heterozygous (9.3 ± 0.4 mmol/l, n = 24, *p* < 0.001) and KO (10.2 ± 0.6 mmol/l, n = 24, *p* < 0.001) mice compared with WT mice (6.2 ± 0.2 mmol/l, n = 24) (Fig. 1B).

**FIGURE 1.**
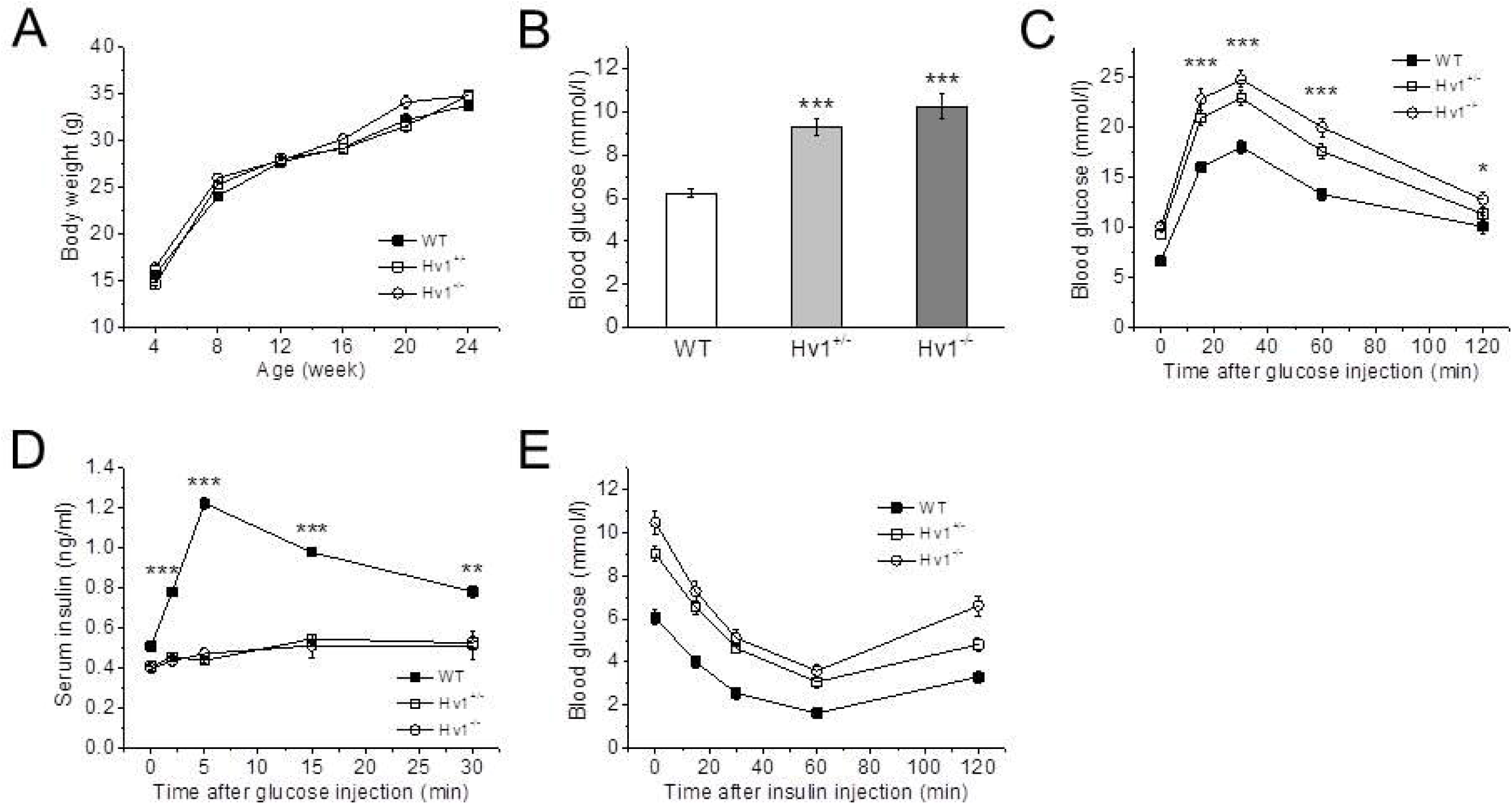
Hv1-deficient mice exhibit hyperglycaemia and impaired glucose tolerance with reduced insulin secretion. A: Body weight change of KO, heterozygous and WT mice (n^+/+^ = 24; n^+/−^ = 24; n^−/−^ = 24). Data are mean ± SEM. B: Basal blood glucose concentrations after fasting 6 h in 4 month-old KO, heterozygous and WT mice (n^+/+^ = 24; n^+/−^ = 24; n^−/−^ = 24). Data are mean ± SEM. *** *p* < 0.001, KO or heterozygous vs. WT. C: Blood glucose levels measured in whole blood following i.p. injection of glucose (2 g/kg body weight) in KO, heterozygous and WT mice (n^+/+^ = 12; n^+/−^ = 12; n^−/−^ = 12). Data are means ± SEM. * *p* < 0.05, \*\*\**p* < 0.001, KO vs. WT. D: Insulin concentrations measured in sera of KO, heterozygous and WT mice following i.p. glucose challenge (2 g/kg body weight) (n^+/+^ = 8; n^+/−^ = 8; n^−/−^ = 8). Data are means ± SEM. \**p* < 0.05, \*\*\**p* < 0.001, KO vs. WT. E: Blood glucose levels measured in whole blood following i.p. insulin injection (1 U/kg body weight) in KO, heterozygous and WT mice (n^+/+^ = 12; n^+/−^ = 12; n^−/−^ = 12). Data are means ± SEM.

To evaluate the impact of Hv1 on disposal of a glucose load, intraperitoneal (i.p.) glucose tolerance tests (IPGTT) were performed. Compared with WT mice, both KO and heterozygous mice in 4 months of age showed significantly higher glucose levels following an i.p. glucose load (2 g/kg body weight) (Fig. 1C). Corresponding serum insulin levels (including basal) were significantly lower in both KO and heterozygous mice throughout the IPGTT after the glucose challenge compared with WT mice, providing evidence for an insulin secretion defect in response to glucose (Fig. 1D). Thus, the Hv1KO mice exhibit an impairment in their ability to dispose of a glucose load due to insulin secretion defect.

To explore the possibility that the observed glucose intolerance was the result of peripheral insulin resistance, we performed i.p. insulin tolerance tests (IPITT) in Hv1KO mice in 4 months of age. We found that insulin administration lowered blood glucose levels in both WT and Hv1-deficient mice to a similar extent, indicating that Hv1 deficiency does not impair a peripheral insulin sensitivity (Fig. 1E). Taken together, these data are compatible with the notion that loss of Hv1 results in impaired glucose tolerance due to a defect of insulin secretion *in vivo*.

### Reduced insulin secretion of islets from Hv1-deficient mice

To delineate the role of Hv1 in insulin secretion, we performed insulin secretion assays using isolated islets from wild type (WT, Hv1^+/+^), heterozygous (Hv1^+/−^) and homozygous (KO, Hv1^−/−^) mice. As shown in Fig. 2A, 16.7 mM glucose-induced insulin secretion was greatly reduced by 51 (n = 8, *p* < 0.001) and 78% (n = 8, *p* < 0.001) in heterozygous and KO islets compared with WT islets (n = 8). While, basal insulin secretion (at 2.8 mM glucose) in heterozygous and KO islets was also significantly reduced by 60 (n = 8, *p* < 0.001) and 80% (n = 8, *p* < 0.001) compared with WT islets (n = 8). Direct depolarization elicited by an increase of extracellular K^+^ (60 mM KCl) also attenuated insulin secretion by 59 (n = 8, *p* < 0.001) and 65% (n = 8, *p* < 0.001) in heterozygous and KO islets compared with WT islets (n = 8) (Fig. 2A), indicating that knockout of Hv1 prevents K^+^-induced insulin secretion in pancreatic *β* cells. Together, these data indicate that loss of Hv1 in islets inhibits insulin secretion.

**FIGURE 2.**
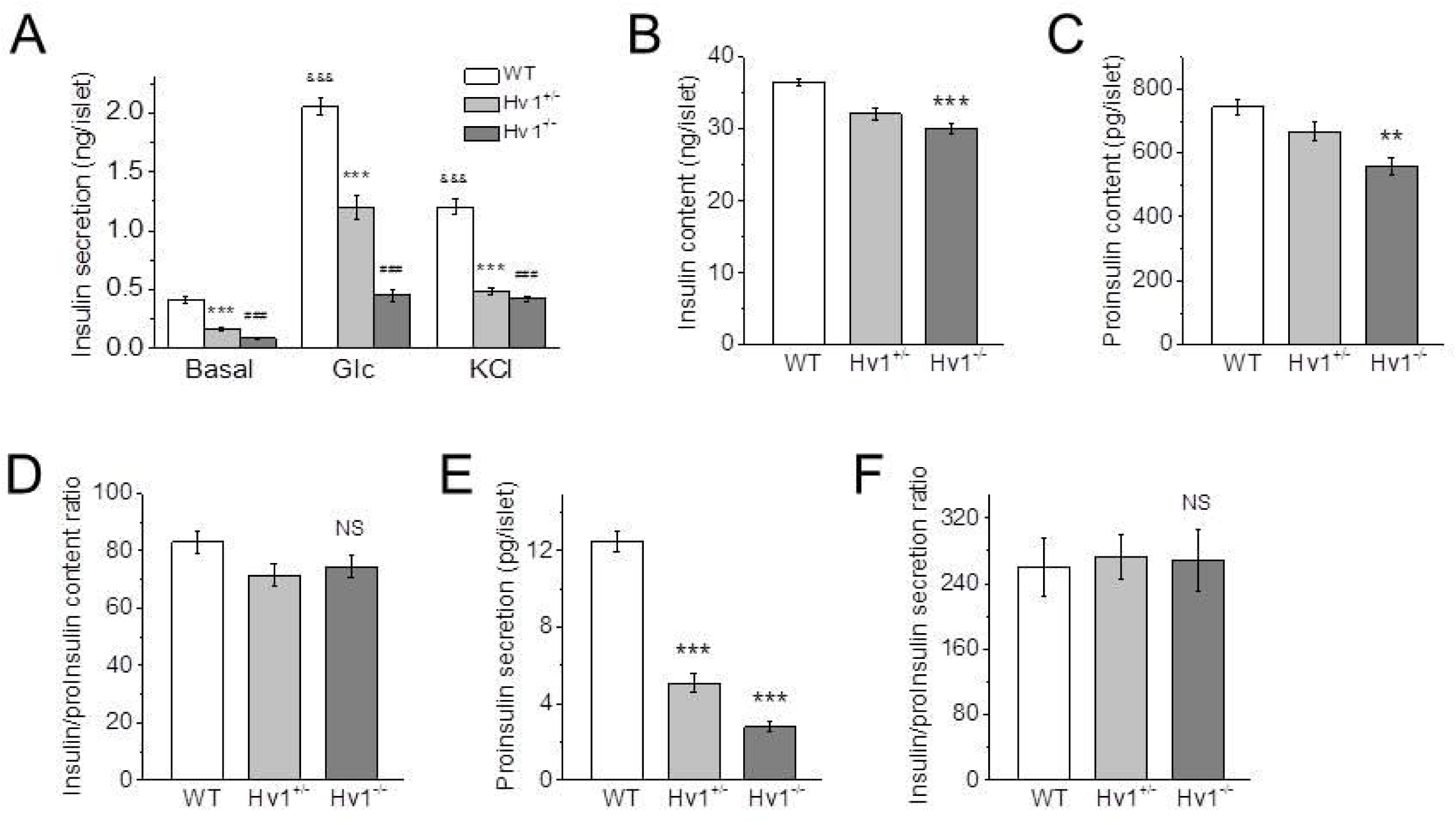
Reduced insulin secretion of islets from Hv1-deficient mice. A: Glucose- and KCl-induced insulin secretion from isolated islets of KO, heterozygous and WT mice (n = 8 per genotype). Data are means ± SEM. *** *p* < 0.001, heterozygous vs. corresponding WT; ^###^*p* < 0.001, KO vs. corresponding WT; ^&&&^*p* < 0.001, Glc or KCl vs. Basal for WT. Basal, 2.8 mM glucose; Glc, 16.7 mM glucose; KCl, 60 mM KCl. B and C: Insulin (B) and proinsulin (C) contents at a basal condition (2.8 mM glucose) in isolated islets from KO, heterozygous and WT mice (n = 8 per genotype). Data are means ± SEM. \*\**p* < 0.01, \*\*\**p* < 0.001, KO vs. corresponding WT. D: Ratio of insulin to proinsulin content of isolated islets of KO, heterozygous and WT mice at a basal condition (n = 8 per genotype). Data are means ± SEM. NS (no significance), vs. WT. E: Proinsulin secretion at 16.7 mM glucose from isolated islets of KO, heterozygous and WT mice (n = 8 per genotype). Data are mean ± SEM. *** *p* < 0.001, KO or heterozygous vs. WT. F: Ratio of insulin to proinsulin secretion of isolated islets of KO, heterozygous and WT mice at 16.7 mM glucose (n = 8 per genotype). Data are means ± SEM. NS (no significance), vs. WT.

Insulin and proinsulin contents in KO islets were reduced by 17 (n = 8, *p* < 0.001) and 25% (n = 8, *p* < 0.01), respectively, compared with WT islets (n = 8) at a basal condition (2.8 mM glucose) (Fig. 2B and C). The ratio of insulin to proinsulin content, however, was indistinguishable between KO and WT islets (Fig. 2D), suggesting that insulin synthesis is abnormal, but not insulin maturation in Hv1-deficient islets. The proinsulin secretion was barely detectable under basal conditions (2.8 mM glucose) in WT, heterozygous and KO islets (data not shown). In the presence of 16.7 mM glucose, the proinsulin secretion was reduced by 59 (n = 8, *p* < 0.001) and 78% (n = 8, *p* < 0.001) in heterozygous and KO islets, compared with WT islets (n = 8) (Fig. 2E). However, the ratios of insulin to proinsulin secretion in heterozygous and KO islets in the presence of 16.7 mM glucose were not different with WT islets (Fig. 2F), suggesting that no significant accumulation of proinsulin occurred in KO islets.

### Knockdown of Hv1 has an effect on insulin synthesis

To further examine the effect of Hv1 on insulin secretion, we used RNA interference to instantaneously reduce endogenous Hv1 levels in INS-1 (832/13) cells. The insulin secretion of INS-1 (832/13) cells at a basal condition (2.8 mM glucose) was low at both the controls and Hv1-knockdown INS-1 (832/13) cells (Fig. 3A). In the presence of 16.7 mM glucose, the insulin secretion in the control INS-1 (832/13) cells increased 9.6-fold compared with that at the basal condition. Whereas the insulin secretion was significantly reduced by 64% (*p* < 0.001) by a reduction in Hv1 level with Hv1-targeting siRNA (Fig. 3A), indicating that Hv1 markedly affects the glucose-induced insulin secretion in INS-1 (832/13) cells.

**FIGURE 3.**
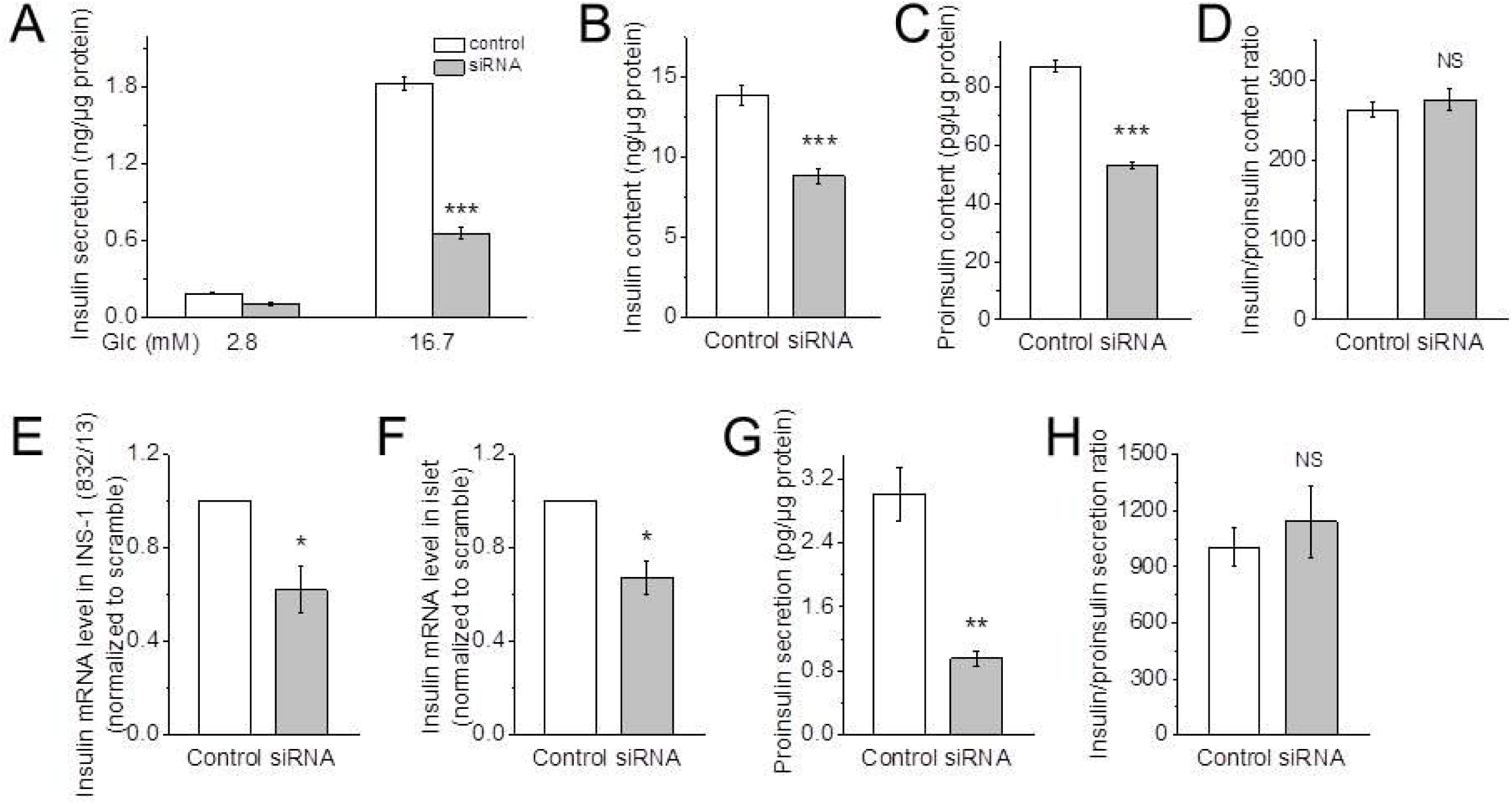
siRNA-mediated knockdown of Hv1 has an effect on insulin synthesis. A: Glucose-induced insulin secretion from INS-1 (832/13) cells transfected with scramble (control) or Hv1-targeting siRNA (siRNA) (n = 8 per condition). Data are mean ± SEM. \*\*\**p* < 0.001, vs. corresponding control. B and C: Insulin (B) and proinsulin (C) contents in the cells transfected with scramble (control) or Hv1-targeting siRNA (siRNA) at a basal condition (n = 8 per condition). Data are mean ± SEM. *** *p* < 0.001, vs. corresponding control. D: The ratio of insulin to proinsulin content of the cells transfected with scramble (control) or Hv1-targeting siRNA (siRNA) at a basal condition (n = 8 per condition). Data are mean ± SEM. NS (no significance), vs. control. E and F: Quantitative RT-PCR for insulin from INS-1 (832/13) cells (E) and isolated islets (F) treated with scramble (control) or Hv1-targeting siRNA (siRNA). Data are mean ± SEM (n = 3 per condition). \**p* < 0.05, vs. corresponding control. G and H: Proinsulin secretion (G) and the ratio of insulin to proinsulin secretion (H) of INS-1 (832/13) cells transfected with scramble (control) or Hv1-targeting siRNA (siRNA) in the presence of 16.7 mM glucose. Data are mean ± SEM (n = 8 per condition). ** *p* < 0.01, \*\*\**p* < 0.001, NS (no significance), vs. corresponding control. These data suggests that no significant accumulation of proinsulin occurred in KO islets.

To examine the effect of Hv1 on insulin processing, the insulin and proinsulin contents were measured. At a basal condition (2.8 mM glucose), the contents of both insulin and proinsulin were reduced by 36 (*p* < 0.001) and 39% (*p* < 0.001) respectively, in the Hv1-silenced INS-1 (832/13) cells, compared with the controls (Fig. 3B and C). However, the ratio of insulin to proinsulin content has a no difference between the controls and the Hv1-downregulated INS-1 (832/13) cells (Fig. 3D), demonstrating that Hv1 has an impact on proinsulin synthesis. To further verify the effect of Hv1 on insulin synthesis, the insulin mRNA expression levels in the INS-1 (832/13) cells and the isolated islets were measured. As shown in Fig 3E and F, insulin mRNA level was decreased by 38 (*p* < 0.01) and 33% (*p* < 0.01) by a reduction in Hv1 level with Hv1-targeting siRNA in the INS-1 (832/13) cells and islets, respectively, compared with the controls, indicating that knockdown of Hv1 has an effect on insulin synthesis.

To detect the effect of Hv1 on proinsulin secretion, we then measured the proinsulin secretion in the INS-1 (832/13) cells. The proinsulin secretion was barely detectable under a basal condition in the control, but in the presence of 16.7 mM glucose, the proinsulin secretion was reduced by 68% (*p* < 0.01) by a reduction in Hv1 level, compared with the control (Fig. 3G). However, the ratio of insulin to proinsulin secretion in Hv1-silenced INS-1 (832/13) cells in the presence of 16.7 mM glucose was not different with that in control INS-1 (832/13) cells (Fig. 3H), suggesting that no significant accumulation of proinsulin occurred in the Hv1-silenced INS-1 (832/13) cells.

### Deficiency of Hv1 reduces *β* cell masses and pancreatic insulin content

To determine whether Hv1 deletion affects islet development, we conducted immunohistologic (IHC) studies. Morphometric analysis of pancreatic sections from WT, heterozygous and KO mice at 4 months of age exhibited a relatively normal islet architecture in each case, with *β* cells concentrated in the core and *α* cells located mainly in the periphery (Fig. 4A), while the morphology of isolated islets cultured overnight from KO mice is not overtly different from WT islets (data not shown). On the other hand, the number of the isolated islets per pancreas is not significantly different between WT and KO mice (data not shown), which is consistent with the result from immunohistochemical analysis (Fig. 4B). However, the islet average size calculated from isolated islets (Fig. 4C) and islet area to total pancreas area (Fig. 4D) analyzed by immunohistochemistry of pancreatic sections were decreased in the KO mice compared with WT and heterozygous mice.

**FIGURE 4.**
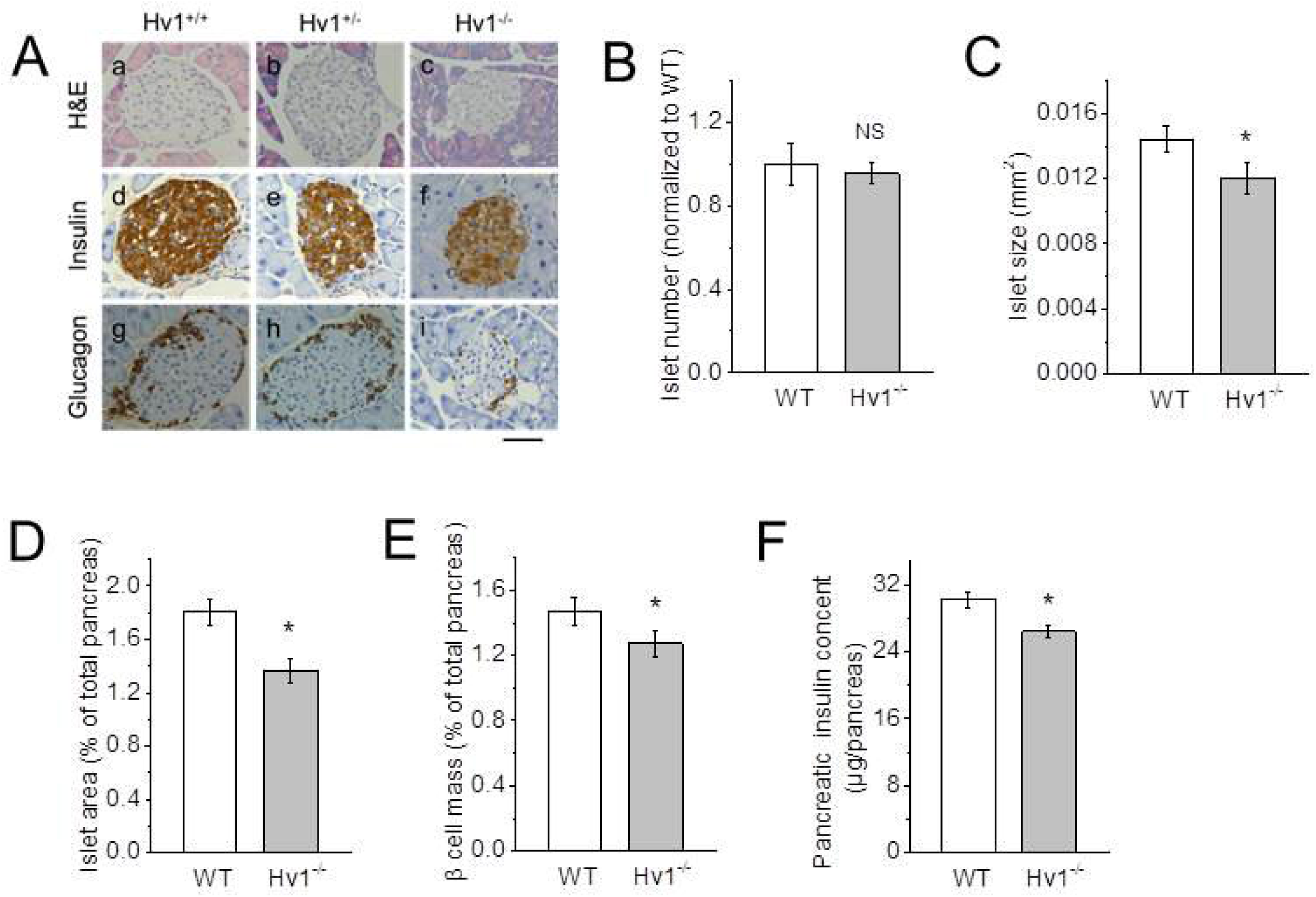
Knockout of Hv1 decreases *β* cell mass. A: Immunohistochemical analysis of 4 month-old WT, heterozygous and KO islets using anti-insulin and anti-glucagon antibodies. Representative images of H&E, anti-insulin and anti-glucagon antibody-stained pancreatic sections from 4 month-old WT, heterozygous and KO mice. Scale bar, 50 μm. B and C: Relative islet number based on immunohistochemical analysis of pancreatic sections (B) and islet size calculated by isolated islet area (C) in 4 month-old WT and Hv1KO mice (n = 6 per genotype). Data are mean ± SEM. NS (no significan ce), \**p* < 0.05, vs. WT. D: Relative islet area of 4 month-old WT and KO mice analyzed by immunohistochemistry of pancreatic sections (n = 6 per genotype). Data are mean ± SEM. * *p* < 0.05, vs. WT. E: *β* cell mass of 4 month-old WT and KO mice based on immunostaining of pancreatic sections (n = 6 per genotype). Relative *β* cell mass was determined as a ratio of total insulin-positive area to total pancreatic area. Twenty to thirty sections per pancreas were analyzed. Data are presented as mean ± SEM. * *p* < 0.05, \*\**p* < 0.01, vs. corresponding WT. F: Pancreatic insulin content (n = 6 per genotype). Data are mean ± SEM. * *p* < 0.05, vs. WT.

The quantification of total *β* cell mass displayed a genotype-dependent difference. *β* cell mass was decreased by 13% (n = 6, *p* < 0.05) in Hv1KO mice compared with WT mice (n = 6), as measured by morphometric analysis of insulin-positive islet cells (Fig. 4E). The total pancreatic insulin content in Hv1KO mice was also decreased by 11% (n = 6, *p* < 0.05) (Fig. 4F), the same as the observed in the isolated islets (Fig. 2B). These results show that Hv1KO mice have sufficient *β* cells and insulin, indicating that the *in vivo* phenotype is not due to gross developmental defects.

### Deficiency of Hv1 affects the size of insulin granules, but not the number and docking of the vesicles

Insulin is stored in large dense-core secretory granules in *β* cells and released via granule exocytosis upon stimulation. The correct size, number and docking to cell membrane of insulin granules are necessary for insulin secretion in *β* cells (19,20). We used TEM to investigate whether loss of Hv1 disturbs vesicle distribution in *β* cells. The ratios of the small size vesicles (<300nm) and large size vesicles (>400nm) were increased by 4-fold and decreased by 47% in Hv1KO mice compared with that in WT mice, respectively, but the ratio of the middle size vesicles (300-400nm) showed no significant difference (Fig. 5A and B). However, the total number of the vesicles in *β* cells in Hv1KO mice is the same with that in WT mice (Fig. 5C). This result might explain why there is a difference of insulin contents between Hv1KO and WT mice (Fig. 1B). The docking of insulin granules plays an important role in regulating insulin secretion (25). The detailed quantitative electron microscopic analysis of *β* cells was performed to study the docked granules. The number of secretory granules close to (<100 nm) the cell membrane in *β* cells in Hv1KO mice is similar to that in WT mice (Fig. 5D), indicating that the loss of Hv1 does not influence the granule docking in *β* cells.

**FIGURE 5.**
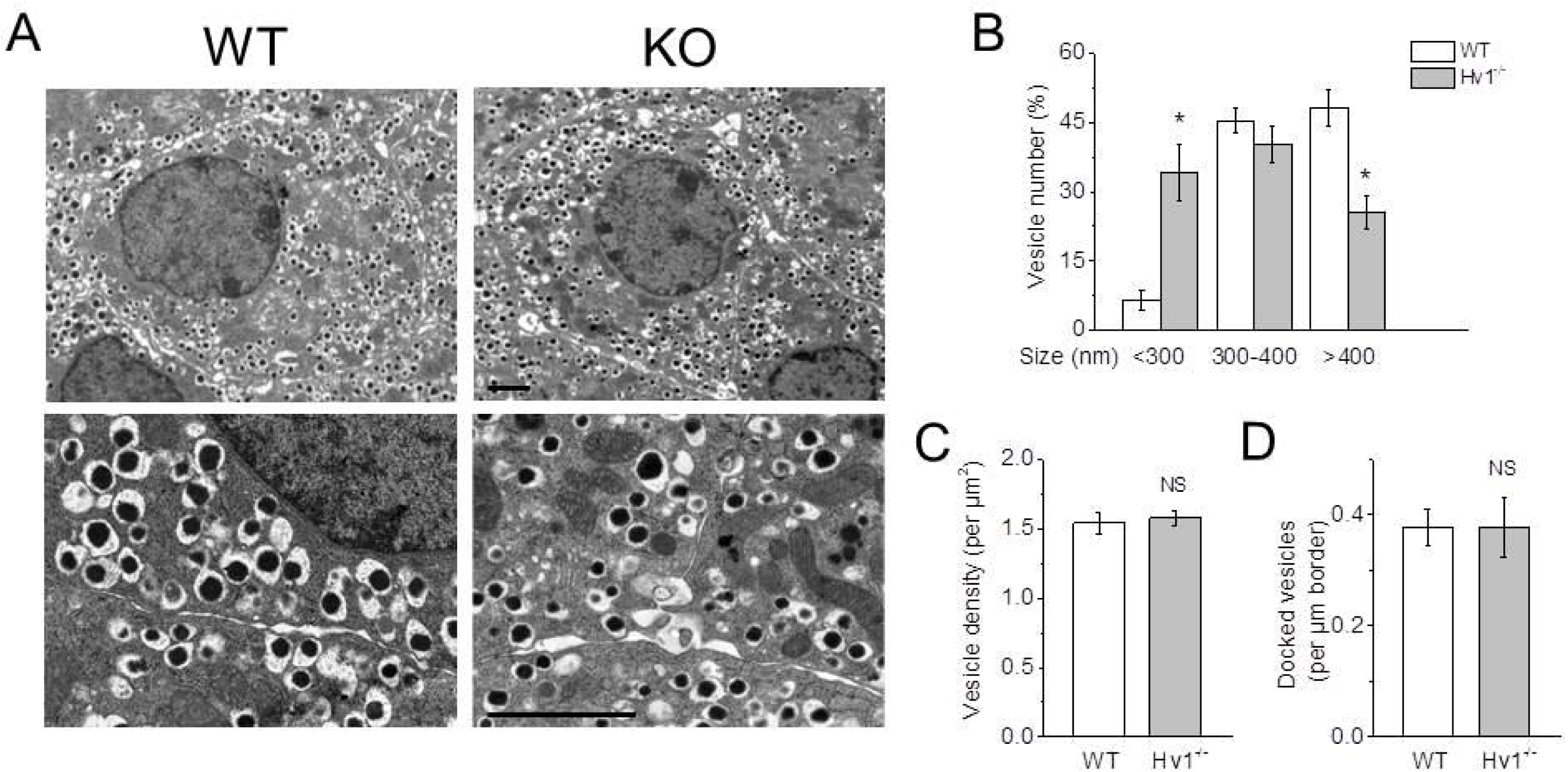
Deficiency of Hv1 affects the size of insulin granules, but not the number and docking of the vesicles. A: Representative TEM images of *β* cells from 4 month-old WT and KO mice. Scale bar, 2 μm. B: Relative size distribution of vesicles in *β* cells based on TEM images. The size of vesicles in Hv1KO mice shows smaller than WT mice (n = 50 per genotype). Data are mean ± SEM. * *p* < 0.05, vs. WT. C: Vesicle density in pancreatic *β* cells of 4 month-old WT and KO mice analyzed by TEM images (n = 50 per genotype). Data are mean ± SEM. NS (no significance), vs. WT. D: Number of vesicles docked onto cytoplasmic membrane in *β* cells of WT and KO islets. from TEM-based analysis (n=50 per genotype). Data are mean ± SEM. NS (no significance), vs. WT.

### Hv1 regulates insulin secretion dependent on PKC activation

cAMP is an important second messenger involved in potentiating rather than initiating insulin secretion (21). To detect the effect of Hv1 on cAMP production, we measured forskolin enhanced insulin secretion and cAMP content in isolated islets. In the present of 10 μM forskolin, at 2.8 mM glucose, the insulin secretion in Hv1-silenced islets was decreased by 37% (*p* < 0.01) compared with that in control islets, while at 16.7 mM glucose, the insulin secretion was reduced by 63% (p < 0.001) (Fig. 6A). The cAMP content in Hv1-silenced islets was not different with that in control islets in the present of 10 μM forskolin and 16.7 mM glucose (Fig. 6B), suggesting that knockdown of Hv1 has no effect on cAMP production.

**FIGURE 6.**
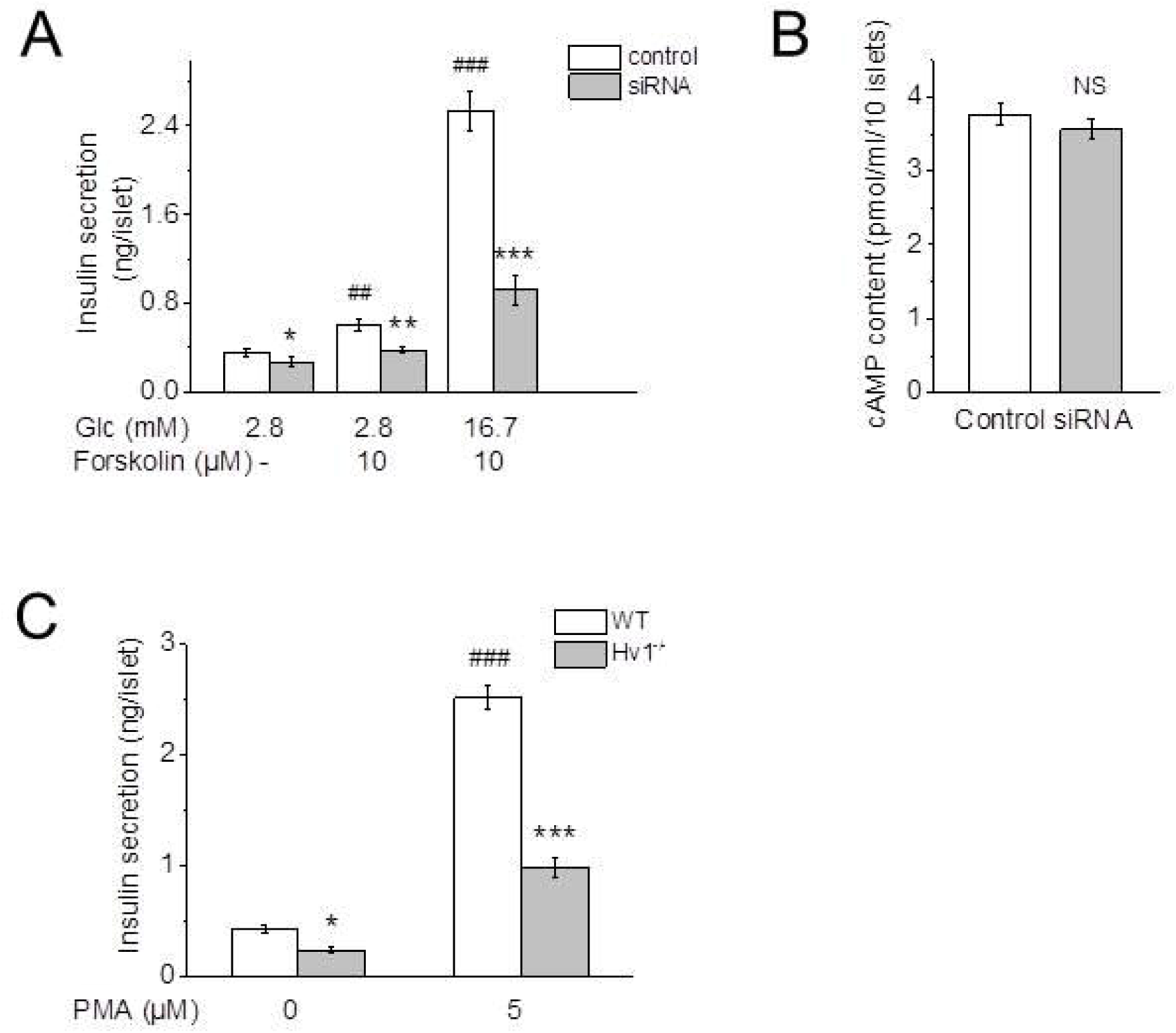
Hv1 regulates insulin secretion through PKC signaling pathway. A: Insulin secretion of isolated islets transfected with scramble (control) or Hv1-targeting siRNA (siRNA) at 2.8 mM glucose, 2.8 mM glucose containing 10 μM forskolin, and 16.7 mM glucose containing 10 μM forskolin (n = 8 per condition). Data are mean ± SE M. \**p* < 0.05, \*\**p* < 0.01, \*\*\**p* < 0.001, vs. corresponding control; ^##^*p* < 0.01, ^###^*p* < 0.001, vs. control at 2.8 mM glucose. B: cAMP contents of isolated islets transfected with scramble (control) or Hv1-targeting siRNA (siRNA) at 16.7 mM glucose containing 10 μM forskolin (n = 6 per condition). Data are mean ± SEM. NS (no significance), vs. control. C: PMA-induced insulin secretion from isolated islets of WT and KO mice (n = 8 per genotype). Data are means ± SEM. \**P* < 0.05, \*\*\**p* < 0.001, vs. corresponding WT; ^###^*p* < 0.001, vs. corresponding at 0 μM PMA.

PMA as a well-known activator of Hv1 has been extensively used for Hv1 function studies (22,23). To examine the effect of Hv1 on protein kinase C (PKC) signaling pathway, we measured PMA-induced insulin secretion in isolated islets from WT and Hv1KO mice. As shown in Fig. 6C, in the presence of 5 μM PMA, the insulin secretion in the isolated islets from WT mice increased 6.3-fold compared with that in the absence of PMA, whereas the insulin secretion in Hv1KO islets was significantly decreased by 61% (*p* < 0.001) compared with that in WT islets. These data indicate that Hv1 regulates insulin secretion dependent on PKC activation.

### Deficiency of Hv1 limits Ca^2+^ entry and membrane depolarization

Glucose stimulates insulin secretion by induction of Ca^2+^-dependent electrical activity that triggers exocytosis of the insulin granules. We found that knockout of Hv1 impaired glucose-induced intracellular Ca^2+^ homeostasis (Fig. 7A and B). In pancreatic *β* cells, the increase of cytosolic Ca^2+^ ([Ca^2+^]c) occurs with Ca^2+^ entry across voltage-sensitive Ca^2+^ channels activated by membrane depolarization (24). To confirm whether the abrogation of Ca^2+^ influx by knockout of Hv1 is coupled to membrane polarization, the membrane potential changes of isolated islet *β* cells during glucose-stimulation were monitored with DiBAC_4_(3) fluorescence. As shown in Fig. 7C, the isolated islet *β* cells from WT mice were depolarized significantly more than isolated islet *β* cells from Hv1KO mice after glucose stimulation, indicating that Hv1 deficiency impairs glucose-induced membrane depolarization. Thus, the reduction in insulin secretion by deficiency of Hv1 should involve in electrical activity and [Ca^2+^]c signaling.

**FIGURE 7.**
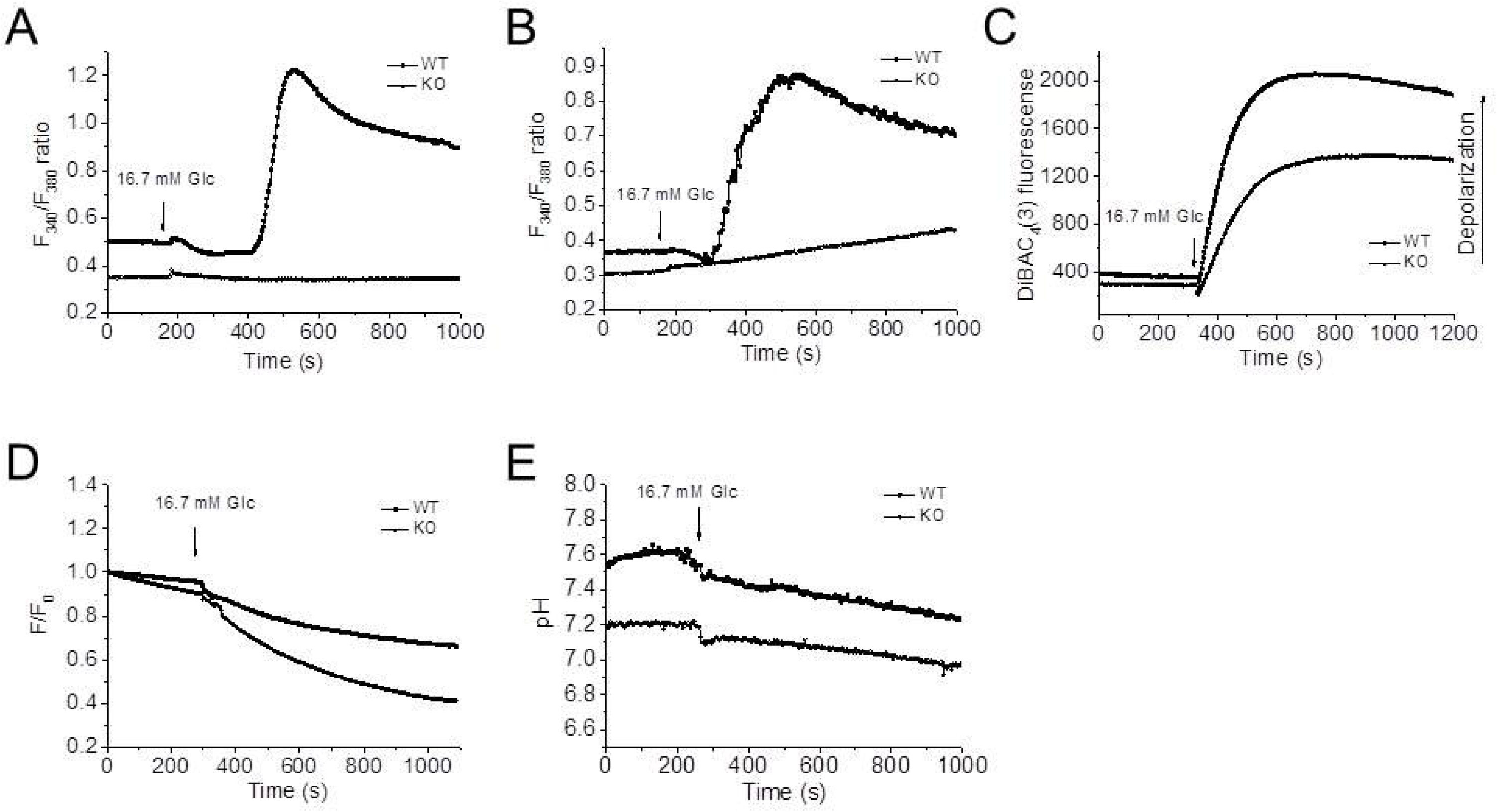
Deficiency of Hv1 limits Ca^2+^ entry and membrane depolarization, decreases cytosolic pH and increases intragranular pH. A and B: Knockdown of Hv1 limits Ca^2+^ entry in isolated islets (A) and isolated islet *β* cells (B) from WT and Hv1KO mice. Cellular Ca^2+^ levels in isolated islets and *β* cells were measured by Fura-2 fluorescence, which were kept at 2.8 mM glucose before switching to stimulation by 16.7 mM glucose as indicated above the traces. C: Deficiency of Hv1 limits membrane depolarization induced by glucose both in isolated islet *β* cells. The membrane potential changes of isolated islet *β* cells during glucose-stimulation were monitored by DiBAC_4_(3) fluorescence. The isolated islet *β* cells from WT and Hv1KO mice were kept at 2.8 mM glucose before switching to stimulation by 16.7 mM glucose as indicated above the traces. D: Knockout of Hv1 results in an increase in intragranular pH monitored by LysoSensor Green DND-189 fluorescence. The data are presented as ratios of LysoSensor Green DND-189 intensity (*F*) over the corresponding initial fluorescence intensity (*F*_0_). E: Deficiency of Hv1 reduces cytosolic pH in isolated islet *β* cells determined by BCECF fluorescence. The arrows indicated the time when the high glucose (final concentration, 16.7 mM) solution was added into the culture dishes, without interruptions in the recordings.

### Knockout of Hv1 alkalinizes intragranules and acidifies cytosol in *β* cells

Our previous results showed that Hv1 is mainly expressed in insulin-containing granules (18). To assess the effect of Hv1 on insulin granule pH, the intragranule pH was monitored by LysoSensor Green DND-189 (LSG) fluorescence. LSG accumulates in acidic organelles and exhibit a pH-dependent fluorescence. LSG fluorescence increases with a pH decrease, but no fluorescence in alkalinization (25). The ratio of LSG fluorescence in Hv1-deleted *β* cells were lower than that of the controls at a basal (2.8 mM glucose) condition, indicating that the intragranules in Hv1-deleted *β* cells were alkalinized (Fig. 7D). While the addition of high glucose solution to *β* cells (final concentration, 16.7 mM glucose), induced the ratio further declined in both Hv1-deleted and control *β* cells. Therefore, these observations indicated that Hv1 involves in intragranule pH regulation in pancreatic *β* cells.

To assess the effect of Hv1 on cytosolic pH in *β* cells, cytosolic pH was evaluated by BCECF fluorescence. As shown in Fig. 7E, the pH in Hv1-knockout *β* cells at both 2.8 and 16.7 mM glucose was lower than that in control *β* cells, indicating that the deficiency of Hv1 in *β* cells decreased cytosolic pH. While stimulation with glucose (final concentration, 16.7 mM glucose) resulted in pH declined in both Hv1-deleted and control *β* cells, indicating that glucose stimulation induces insulin granule acidification.

## DISCUSSION

Here, we show *in vivo*, that both Hv1KO and heterozygous mice display hyperglycemia and glucose intolerance due to markedly decreased insulin secretion. *In vitro*, deficiency of Hv1 exhibits a remarkable defect in glucose-induced insulin secretion, decreases both insulin and proinsulin contents, impairs intracellular Ca^2+^ homeostasis and membrane depolarization, alkalinizes secretory granules and acidifies cytosol. These data indicate that the level of Hv1 expression in the *β* cell is required for insulin secretion and maintaining glucose homeostasis, and reveal a significant role for the proton channel in the modulation of pancreatic *β* cell function.

The acidification inside the secretory vesicles has a fundamental importance for maintenance of whole-body homeostasis. It has been suggested for decades that the proton gradients might be involved in the fusion of secretory vesicles to the target membrane (26,27). Barg et al. (15) showed that the acidic pH might regulate priming of the granules for secretion, a process involving pairing of SNARE proteins on the vesicles and target membranes to establish fusion competence. The dependence of the insulin secretion on Hv1 activity reflects that the intragranule pH is directly involved in the secretion of secretory granules. We presumed that loss of Hv1 would result in abnormal secretory vesicle alkalinization, and thereby affect insulin synthesis.

The effects of glucose on the cytosolic pH (pH_c_) of the islet *β* cells are debated (4-8,28,29). More confusing is the coupling between the pH_c_ changes and insulin secretion. Studies on changes in *β* cell pH_c_ by manipulation of extracellular pH and ionic composition proposed that alkalinization inhibited insulin secretion (3,9), while acidification increased insulin release (10). Although NHE1 and NHE7 are the most expressed in mouse islets, upon stimulation by glucose, the pH_c_ similarly increased in NHE1 mutant and control islets (30), suggesting that β cell alkalinization by glucose is not mediated by Na^+^/H^+^ exchangers as sometimes proposed (6,28). However, either a decrease or an increase in pH_c_ is important for glucose-induced insulin secretion via the triggering or the amplifying pathways. Manipulations of extra- and intracellular pH in pancreatic islets or *β* cells are associated with changes in membrane potential, ionic fluxes, and insulin release (1,3). Glucose-induced priming of insulin secretion, which is thought to be mediated by the amplifying pathway (31), has been proposed to link to the pH changes in *β* cells (32).

As a well-known PKC activator,PMA can activate PKC as an analog of DAG, and the activation of PKC results in membrane depolarization, Ca^2+^ influx and then priming of insulin secretion (33). PMA, as a well-known Hv1 activator (16,17), induces insulin secretion through activating Hv1 activity, suggesting that Hv1 regulates insulin secretion through the PKC signaling pathway. This important defect that ablates the Hv1 gene could be predicted from the reduced depolarization as we observed in the *β* cell. This indicates that *β* cells lacking Hv1 proton channel cannot generate Ca^2+^ influx when membrane depolarization is limited.

The fact that heterozygous mice also have a hyperglycemia with a low insulin level illustrates that Hv1 is at an important control point in the metabolic pathway regulating insulin secretion, and that relatively small changes in Hv1 activity are likely to have important effects on insulin secretion. Similar effects have been observed for glucokinase (34). These data further confirms that the Hv1 is closely related to insulin secretion. The findings of the present study clearly demonstrate that Hv1 plays an important role in positively regulating glucose-stimulated insulin secretion.

*β* cell failure is associated with not only decreased *β* cell insulin secretory function but also reduced overall *β* cell mass (35). In present study, there was no difference in islet morphology between Hv1KO and WT mice, and only a very modest decrease in islet size and *β* cell mass. The smaller size observed in Hv1-deficient pancreatic islets may be related to the decrease in *β* cell mass, and result from impaired insulin secretory function. In this context, it is important to note that *in vitro* siRNA-mediated knockdown of Hv1 in isolated islets and INS-1 (832/13) cells caused decreased glucose-stimulated insulin secretion (GSIS), suggesting that the *in vivo* decrease in insulin secretion in the Hv1KO mice was not due to an *in vivo β* cell developmental defect.

## EXPERIMENTAL PROCEDURES

Additional experimental procedures are provided in *SI Methods*.

## Acknowledgements

We are deeply grateful to Dr. Claes B. Wollheim and Dr. Nicolas Demaurex (Department of Cell Physiology and Metabolism, University of Geneva) for their reviewed the manuscript and many helpful discussions about the manuscript. We would like to thank Dr. Y. Okamura (School of Medicine, Osaka University) for providing VSOP/Hv1 KO mice. We would like to thank Dr. Hans E. Hohmeier (Duke University Medical Center) for providing materials mentioned in the text. This work was supported by National Natural Science Foundation of China (No. 31271464).

## Author contributions

SJL conceived and designed the study. HMP, XDW, WX, QZ, SRZ, JWQ, JLL, YZC, WYZ and SJL performed the experiments. SJL and HMP wrote the paper. SJL, HMP and XDW reviewed and edited the manuscript. All authors were involved in data analysis, read and approved the manuscript.

## References

1. Pace, C.S., J.T. Tarvin, and J.S. Smith. Stimulus-secretion coupling in beta-cells: modulation by pH. Am. J. Physiol. 244: E3–18 (1983).

2. Lindström, P., and J. Sehlin. Effect of glucose on the intracellular pH of pancreatic islet cells. Biochem. J. 218:887–892 (1984).

3. Lindström P., and J. Sehlin. Effect of intracellular alkalinization on pancreatic islet calcium uptake and insulin secretion. Biochem. J. 239:199–204 (1986).

4. Hellman, B., J. Sehlin, and I.B. Taljedal. The intiacellular pH of mammalian pancreatic beta-cells. Endocrinology 90:335-337 (1972).

5. Grapengiesser, E., E. Gylfe, and B. Hellman. Regulation of pH in individual pancreatic ß-cells as evaluated by fluorescence ratio microscopy. Biochim. Biophys. Acta 1014:219–224 (1989).

6. Juntti-Berggren, L., P. Arkhammar, T. Nilsson, P. Rorsman, and P.O. Berggren. Glucose-induced increase in cytoplasmic pH in pancreatic beta-cells is mediated by Na^+^/H^+^ exchange, an effect not dependent on protein kinase C. J. Biol. Chem. 266:23537–23541 (1991).

7. Shepherd, R.M., and J.C. Henquin. The role of metabolism, cytoplasmic Ca2^+^, and pH-regulating exchangers in glucose-induced rise of cytoplasmic pH in normal mouse pancreatic islets. J. Biol. Chem. 270:7915–7921 (1995).

8. Salgado, A., A.M. Silva, R.M. Santos, and L.M. Rosario. Multiphasic action of glucose and α-ketoisocaproic acid on the cytosolic pH of Pancreatic β-Cells evidence for an acidification pathway linked to the stimulation of Ca2+ influx. J. Biol. Chem. 271:8738–8746 (1996).

9. Nabe, K., S. Fujimoto, M. Shimodahira, R. Kominato, Y. Nishi, S. Funakoshi, E. Mukai, Y. Yamada, Y. Seino, and N. Inagaki. Diphenylhydantoin suppresses glucose-induced insulin release by decreasing cytoplasmic H^+^ concentration in pancreatic islets. Endocrinology 147:2717–2727 (2006).

10. Pace, C.S., and J.T. Tarvin. pH modulation of glucose-induced electrical activity in B-cells: involvement of Na/H and HCO3/Cl antiporters. J. Membr. Biol. 73:39–49 (1983).

11. Orci, L., M. Ravazzola, M. Amherdt, O. Madsen, A. Perrelet, J.D. Vassalli, and R.G. Anderson. Conversion of proinsulin to insulin occurs coordinately with acidification of maturing secretory vesicles. J. Cell Biol. 103:2273–2281 (1986).

12. Hutton, J.C. The insulin secretory granule. Diabetologia 32:271–281 (1989).

13. Tompkins, L.S., K.D. Nullmeyer, S.M. Murphy, C.S. Weber, and R.M. Lynch. Regulation of secretory granule pH in insulin-secreting cells. Am. J. Physiol. 283:C429–C437 (2002).

14. Eto, K., T. Yamashita, K. Hirose, Y. Tsubamoto, E.K. Ainscow, G.A. Rutter, S. Kimura, M. Noda, M. Iino, and T. Kadowaki. Glucose metabolism and glutamate analog acutely alkalinize pH of insulin secretory vesicles of pancreatic β-cells. Am. J. Physiol. 285:E262–E271 (2003).

15. Barg, S., P. Huang, L. Eliasson, D.J. Nelson, S. Obermuller, P. Rorsman, F. Thevenod, and E. Renstrom. Priming of insulin granules for exocytosis by granular Cl- uptake and acidification. J. Cell Sci. 114, 2145–2145 (2001).

16. Ramsey, I.S., M.M. Moran, J.A. Chong, and D.E. Clapham. A voltage-gated proton-selective channel lacking the pore domain. Nature 440:1213–1216 (2006).

17. Sasaki, M., M. Takagi, and Y. Okamura. A voltage sensor-domain protein is a voltage-gated proton channel. Science 312:589–592 (2006).

18. Zhao, Q., Y. Che, Q. Li, S. Zhang, Y.T. Gao, Y. Wang, X. Wang, W. Xi, W. Zuo, and S.J. Li. The voltage-gated proton channel Hv1 is expressed in pancreatic islet β-cells and regulates insulin secretion. Biochem. Biophys. Res. Commun. 468:746–751 (2015).

19. Lang, J. Molecular mechanisms and regulation of insulin exocytosis as a paradigm of “https://www.ncbi.nlm.nih.gov/pubmed/9914469” endocrine secretion. Eur. J. Biochem. 259:3–17 (1999).

20. Zhao, A., Ohara-Imaizumi, M., Brissova, M., Benninger, R.K., Xu, Y., Hao, Y., Abramowitz, J., Boulay, G., Powers, A.C., Piston, D., Jiang, M., Nagamatsu, S., Birnbaumer, L., and Gu, G. Gαo represses insulin secretion by reducing vesicular docking in pancreatic beta-cells. Diabetes. 59:2522–2529 (2010).

21. Prentki, M., and Matschinsky, F.M. Ca2+, cAMP, and phospholipid-derived messengers in coupling mechanisms of insulin secretion. Physiol. Rev. 67:1185–1248 (1987).

22. Ramsey, I.S., E. Ruchti, J.S. Kaczmarek, and D.E. Clapham. Hv1 proton channels are required for high-level NADPH oxidase-dependent superoxide production during the phagocyte respiratory burst. Proc. Natl. Acad. Sci. U.S.A. 106:7642–7647 (2009).

23. Decoursey, T.E. Voltage-gated proton channels and other proton transfer pathways. Physiol. Rev. 83:475–579 (2003).

24. Rorsman, P., and E. Renstrom. Insulin granule dynamics in pancreatic beta cells. Diabetologia 46:1029–1045 (2003).

25. Renström, E., Eliasson, L., Bokvist, K., and Rorsman, P. Cooling inhibits exocytosis in single mouse pancreatic B-cells by suppression of granule mobilization. J. Physiol. (Lond.) 494:41–52 (1996).

26. Al-Awqati, Q. Proton-translocating ATPases. Annu. Rev. Cell Biol. 2:179–199 (1986).

27. Burgess, T.L., and Kelly, R.B. Constitutive and regulated secretion of proteins. Annu. Rev. Cell Biol. 3:243–293 (1987).

28. Lebrun, P., E.V. Ganse, M. Juvent, M. Deleers, and A. Herchuelz. Na^+^-H^+^ exchange in the process of glucose-induced insulin release from the pancreatic B-cell. Effects of amiloride on 86Rb,45Ca fluxes and insulin release. Biochim. Biophys. Acta 886:448–456 (1986).

29. Lynch, A.M., J.E. Meats, L. Best, and S. Tomlinson. Effect of extracellular sodium removal upon 86Rb outflow from pancreatic islet cells. Biochim. Biophys. Acta 1012:166–170 (1989).

30. Stiernet, P., M. Nenquin, P. Moulin, J.C. Jonas, and J.C. Henquin. Glucose-induced cytosolic pH changes in beta-cells and insulin secretion are not causally related: studies in islets lacking the Na^+^/H^+^ exchanger NHE1. J. Biol. Chem. 282:24538–24546 (2007).

31. Taguchi, N., T. Aizawa, Y. Sato, F. Ishihara, and K. Hashizume. 1995. Mechanism of glucose-induced biphasic insulin release: physiological role of adenosine triphosphate-sensitive K+ channel-independent glucose action. Endocrinology 136:3942–3948 (1995).

32. Gunawardana, S.C., and G.W. Sharp. 2002. Intracellular pH plays a critical role in glucose-induced time-dependent potentiation of insulin release in rat islets. Diabetes 51:105–113 (2002).

33. Shigeto, M., Ramracheya, R., Tarasov, A.I., Cha, C.Y., Chibalina, M.V., Hastoy, B., Philippaert, K., Reinbothe, T., Rorsman, N., Salehi, A., Sones, W.R., Vergari, E., Weston, C., Gorelik, J., Katsura, M., Nikolaev, V.O., Vennekens, R., Zaccolo, M., Galione, A., Johnson, P.R., Kaku, K., Ladds, G., and Rorsman, P. GLP-1 stimulates “https://www.ncbi.nlm.nih.gov/pubmed/26571400” insulin secretion by PKC-dependent TRPM4 and TRPM5 activation. J. Clin. Invest. 125:4714–4728 (2015).

34. Matschinsky, F.M., Glaser, B., and Magnuson, M.A. Pancreatic beta-cell glucokinase: closing the gap between theoretical concepts and experimental realities. Diabetes 47:307–315 (1998).

35. Weir, G.C., and Bonner-Weir, S. Five stages of evolving beta-cell dysfunction during progression to diabetes. Diabetes 53 (Suppl 3):S16–S21 (2004).

